# Misfolded proteins share a common capacity in disrupting LLPS organizing membrane-less organelles

**DOI:** 10.1101/317347

**Authors:** Jian Kang, Liangzhong Lim, Jianxing Song

## Abstract

Profilin-1 mutants cause ALS by gain of toxicity but the underlying mechanism remains unknown. Here we showed that three PFN1 mutants have differential capacity in disrupting dynamics of FUS liquid droplets underlying the formation of stress granules (SGs). Subsequently we extensively characterized conformations, dynamics and hydrodynamic properties of C71G-PFN1, FUS droplets and their interaction by NMR spectroscopy. C71G-PFN1 co-exists between the folded (55.2%) and unfolded (44.8%) states undergoing exchanges at 11.7 Hz, while its unfolded state non-specifically interacts with FUS droplets. Results together lead to a model for dynamic droplets to recruit misfolded proteins, which functions seemingly at great cost: simple accumulation of misfolded proteins within liquid droplets is sufficient to reduce their dynamics. Further aggregation of misfolded proteins within droplets might irreversibly disrupt/destroy structures and dynamics of droplets, as increasingly observed on SGs, an emerging target for various neurodegenerative diseases. Therefore, our study implies that other misfolded proteins might also share the capacity in disrupting LLPS.

---

Amyotrophic lateral sclerosis (ALS) is the most common motor neuron disease, which was first described in 1869 but its mechanism still remains a great mystery (1). Human superoxide dismutases 1 (SOD1) was the first gene identified to be associated with familial ALS and so far more than 20 ALS-causative genes have been found (1-3). Like all other neurodegenerative diseases (4), ALS is also characterized by severe aggregation of these proteins *in vivo* and *in vitro* (1-9). The ALS-associated proteins can be categorized into two major groups: the first including SOD1 and profilin-1 (PFN1) whose wild types are well-folded and only the ALS-associated mutants become aggregation-prone and toxic (1-3,7-10); while the second including FUS and TDP-43 containing the low-complexity (LC) domains whose wild types are already aggregation-prone and toxic (1, 10-15). Very intriguingly, some ALS-causing mutants of the first group such as C71G-PFN1 have been previously revealed to trigger seed-dependent co-aggregation with the proteins in the second group including TDP-43 to manifest the prion-like propagandation (16), but so far the mechanism underlying this observation remains completely unknown.

Progressive aggregation of thousands of non-specific proteins has also been shown to be associated with aging down to unicellular organisms (17,18). Although it remains largely elusive whether the aggregation of non-specific proteins also leads to aging by gain of toxicity, emerging evidence implies that to clean up aggregated proteins represents a key step for cells to rejuvenate as evidenced by the previous result that upon division of *E. coli* cells, the mother cell keeps protein aggregates so as to allow the daughter cell to be free of protein aggregates for rejuvenation (18); as well as by a recent report that for *Caenorhabditis elegans*, upon fertilization the sperm will send a signal to the egg to rejuvenate itself by clearing out protein aggregates via a lysosome-based degradation pathway typically in thirty minutes (19). In this context, protein quality control (PQC) machineries play a central role in handling protein aggregates in cells. Very recently, a novel machinery constituted by stress granules (SGs) and components of the classic PQC machinery has been identified to recruit misfolded proteins for further disaggregation with the assistance of molecular chaperones, or for autophagy-mediated clearance (20-22). SGs are membrane-less compartments formed to respond cellular stress by RNA-binding proteins (RBPs) including FUS and TDP-43 (1,10-14,20-24). Recent studies revealed that SGs are not classically defined complexes such as ribosome, but are dynamic macromolecular assemblies behaving as liquid droplets, whose formation are driven by weak and multivalent interactions among components via a process known as liquid-liquid phase separation (LLPS) (1,10-14,20-27). Remarkably, LLPS has been now recognized to be a general organizing principle for forming membrane-less intracellular compartments/organelles which include nucleolus, Cajal bodies and nuclear speckles in the nucleoplasm, as well as stress granules, P-bodies and germ granules in the cytoplasm (1,27). The self-assembly of dynamic SGs has been now identified to rely on the LC domains of RBPs particularly the prion-like domains of FUS and TDP-43 (1,12-14,20-27). On the other hand, the RBPs of SGs have been also identified to form pathological aggregates in neurons of patients of a variety of neurodegenerative diseases. In particular, the key SG components FUS and TDP-43 have been found in pathological inclusions of ALS and FTD, which led to the hypothesis that pathological inclusions of RBPs are exaggerated from dynamic and functional SGs (1,12-16,20-27).

On the other hand, very recently it has been shown that misfolded proteins, including ALS-associated variants of the high-complexity proteins, have a general tendency to become accumulated and aggregated within SGs (20-22). The accumulation of these proteins in SGs led to aberrant SGs with reduced dynamics of liquid droplets on the one hand, but also promotes the recruitment of chaperones including chaperone HSP70 for SG disassembly on the other hand. However, if the disassembly failed, the aberrant SGs would be transported to the aggresome for degradation by autophagy. Therefore, SG represents a novel surveillance machinery that functions to non-classically recruit misfolded proteins (20-22). This also implies that the misfolded disease-causing variants of the well-folded proteins might gain their toxicity to initiate neurodegenerative diseases including ALS by disrupting dynamics of liquid droplets of SGs. Indeed, SGs have been recently emerging as a central target for various neurodegenerative diseases including ALS (1,10-15,20-29).

So far, the biophysical basis for the liquid droplets formed by the prion-like domains to recruit misfolded proteins remains completely unexplored. Here with screening of the interactions of the FUS/TDP-43 prion-like domains with ALS-causing SOD1/profilin-1 mutants, we have identified C71G-PFN1 and FUS (1-267) to constitute a system suitable for high-resolution characterizations of their solution conformations, dynamics on different time scales and hydrodynamic properties by NMR spectroscopy. The obtain results together allow the proposal of a model for liquid droplets to recruit misfolded proteins, which also appears to underlie gain of toxicity not only for ALS-causing profilin mutants, but also for other misfolded proteins to initiate diseases or even aging by disrupting various cellular membrane-less organelles/compartments formed through LLPS.

## Results

### Identification of a system suitable for high-resolution NMR characterizations

In the present study, we focused on investigating conformations, dynamics, hydrodynamic properties and interaction of ALS-causing PFN1 mutants with liquid droplets of FUS mainly by NMR spectroscopy, because NMR has two significant advantages: 1) NMR requires no introduction of extra tags and chemical groups; and 2) it can offer atomic resolution conformational and dynamic information. This first advantage is particularly critical for the current study because LLPS is extremely sensitive to even minor changes of protein side chains. For example, it has been previously demonstrated that a single-residue mutation was sufficient to alter or even disrupt LLPS of the FUS and TDP-43 LC domains (11-14,23-26). As such, the labelling of extra tags/chemicals required for other methods, such as Fluorescence Recovery after Photobleaching (FRAP), will unavoidably introduce unpredictable perturbations to LLPS or/and interaction between misfolded proteins and liquid droplets.

On the other hand, here one major challenge to perform high-resolution NMR conformational and dynamic studies is to obtain a stable system composed of two proteins: the LC domain of SG components which is able to form dynamic liquid droplets with sufficient number and size; and a misfolded mutant of a well-folded protein which can stably give high-quality NMR spectra. Therefore for the LC domains, we have tested TDP-43 C-terminal LC domain over residues 263-414, FUS N-terminal 1-165 and 1-267 we previously characterized (15,28). Unfortunately, we found that the TDP-43 C-terminal LC domain has two disadvantages: 1) it only formed a small number of small liquid droplets at 25 °C exactly as previously observed (26,30); 2) most likely due to the presence of a hydrophobic fragment 311-341 (15,26), visible aggregates formed upon adding ALS-causing mutants of SOD1 or PFN1. For FUS (1-165) containing only the GQSY-rich prion-like domain, it only formed small liquid droplets at 25 °C as previously reported (25), while FUS (1-267), the entire N-terminal LC domain of FUS with an extra Gly-rich region 166-267 could form dynamic droplets with large numbers and sizes at 25 °C (Fig. 1A and Video 1A).

**Figure 1.**
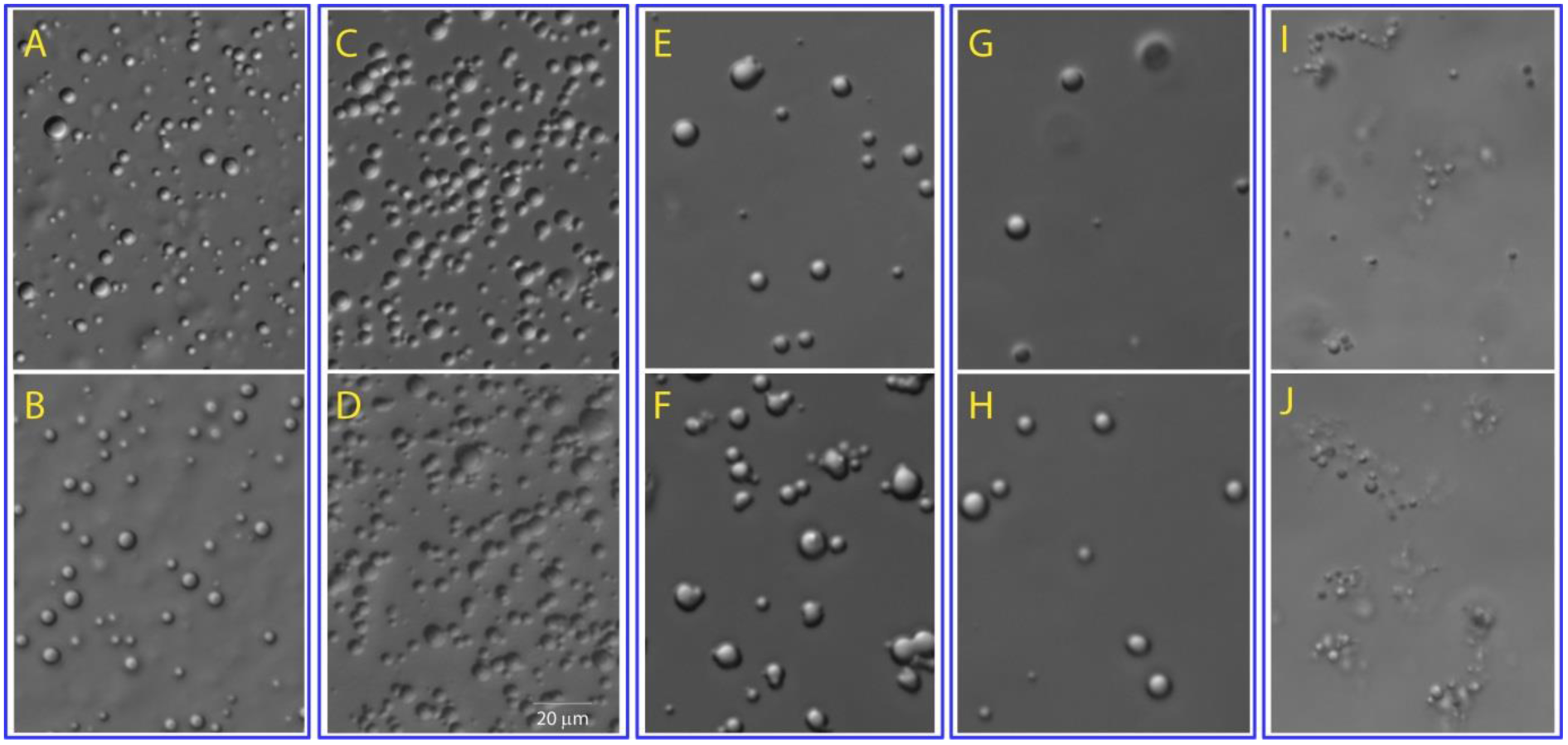
ALS-causing PFN1 mutants disrupt dynamics of liquid droplets. DIC microscopy images of liquid droplets of FUS (1-267) at a protein concentration of 50μM (A); FUS (1-267) + WT-PFN1 at a molar ratio of 1:1 (FUS:PFN1) (B); FUS (1-267) + C71G-PFN1 at a molar ratio of 1:0.5 (C) and at 1:1 (FUS:C71G) (D); FUS (1-267) + G118V-PFN1 at a molar ratio of 1:1 (E) and at 1:2 (FUS:G118V) (F); FUS (1-267) + E117G-PFN1 at a molar ratio of 1:1 (G) and at 1:2 (FUS:E117G) (H); and FUS (1-267) + TDP-43 (10-102) at a molar ratio of 1:0.1 (I) and at 1:0.5 (FUS:TDP-43) (J). The videos for outputting these images are provided in Supplementary Materials.

On the other hand, for the misfolded ALS-causing mutants, we have also tested L126Z-SOD1 with the last 28 residues of SOD1 deleted (7), and ALS-causing PFN1 mutants including C71G-, G118V- and E117G-PFN1. Unfortunately, the solubility of L126Z-SOD1 was very low in buffer and most of its HSQC peaks become too broad to be detectable although no visible aggregates could be seen at low protein concentrations immediately. For PFN1 mutants, our preliminary NMR HSQC characterization showed that both C71G- (Fig. S1A) and G118V-PFN1 (Fig. S1B) have the co-existence of the folded and unfolded states but C71G-PFN1 has the unfolded population much higher than G118V-PFN1. Interestingly, under the same conditions, only the HSQC peaks of the folded state could be detected for E117G-PFN1 (Fig. S1C). In consideration of the previous reports that C71G-PFN1 was the most toxic PFN1 mutant (3,6,8), which has been also recently demonstrated by cell and animal models to trigger ALS by gain of toxicity (6), as well as our observation that C71G-PFN1 could also stably exist in buffer at 25°C to give high-quality NMR spectra for both folded and unfolded states, we thus selected the system composed of FUS (1-267) and C71G-PFN1 for further high-resolution NMR characterizations.

### ALS-causing PFN1 mutants disrupt dynamics of liquid droplets of FUS (1-267)

Previously we have characterized FUS (1-267) to be intrinsically disordered (28). Here we found that it was able to form liquid droplets much larger than FUS (1-165). The diameters of some droplets could even reach ~10 μm at a protein concentration of 50 μM, but they still remain dynamic and undergo Brownian motion in solution as imaged by differential interference contrast (DIC) microscopy (Fig. 1A and Video 1A). On the other hand, as expected, WT-PFN1, C71G-, G118V- and E117G-PFN1 samples were all homogenous and transparent solutions with no droplets detected by DIC microscopy. Interestingly, addition of WT-PFN1 to the FUS (1-267) sample at a protein concentration of 50 μM triggers no significant change of the droplet dynamics at molar ratios of both 1:1 (Fig. 1B and Video 1B1), and at 1:2 (FUS:PFN1) (Video 1B2).

By a sharp contrast, addition of C71G-PFN1 to a molar ratio of 1:0.5 (FUS:C71G) obviously reduced dynamics of droplets and also appeared to trigger the clustering/coalescing of droplets (Fig. 1C and Video 1C). Remarkably, addition of C71G-PFN1 to 1:1 completely disabled Brownian motion of the large droplets (Fig. 1D and Video 1D). Further addition of C71G-PFN1 led to the formation of a gel-like state with high viscosity which gave no NMR signal. Interestingly, addition of G118V-PFN1 with the ALS-causing toxicity lower than C71G-PFN1 (3,6,8) to a molar ratio of 1:1 (FUS:G118V) appeared to only slightly enhance the coalescing of small droplets into larger ones (Fig. 1E and Video 1E). However, addition of G118V-PFN1 to 1:2 was able to deform the round shape and to disrupt Brownian motion of the large droplets (Fig. 1F and Video 1F). On the other hand, addition of E117G-PFN1 with the ALS-causing toxicity the weakest (3,6,8) appeared to only enhance the coalescing of small droplets into larger ones but to induce no significant alteration of the dynamics of liquid droplets at molar ratios (FUS:E117G) of 1:1 (Fig. 1G and Video 1G) and 1:2 (Fig. 1H and Video 1H).

To assess whether other misfolded/unfolded and aggregation-prone proteins also have the capacity in disrupting liquid droplets observed for PFN1 mutants, we further evaluated the effect of TDP-43 (10-102) which is not associated with any diseases. Previously we determined the first 80 residues of TDP-43 (1-102) to adopt a novel ubiquitin-like fold and showed that the deletion the first 9 residues forming the first β-strand of the ubiquitin-like fold resulted in a predominantly disordered TDP-43 (10-102), which is however soluble and stable in buffer at low concentrations (< 100 μM) for several days (31). Remarkably, addition of TDP-43 (10-102) only to a molar ratio of 1:0.1 (FUS:TDP-43) appeared to be sufficient to disrupt the large droplets into small ones, and most interestingly many droplets were connected in the “beads-in-a-string” style (Fig. 1I and Video 1I). Although further addition of TDP-43 (10-102) to 1:0.5 triggered no significant difference immediately (Fig. 1J and Video 1J), the sample formed visible aggregates after ~half an hour.

### NMR characterizations of C71G-PFN1 conformations and dynamic

Profilin-1 is a 140-residue actin-binding protein adopting a well-structured globular fold (Fig. 2), while C71G-PFN1 is the most toxic ALS-causing mutant. Previously we only conducted a preliminary HSQC characterization showing that the C71G mutation rendered PFN1 to co-exist in the folded and unfolded states (9). As such, due to significantly different transverse relaxation time T2 of the folded and unfolded state, even the populations of two states could not be derived from HSQC peak intensity of the co-existing two states as we specially clarified in the previous report (9). Furthermore, other quantitative NMR conformational and dynamic parameters of the co-existing two states, which on the one hand are critical for elucidating the biophysical mechanism for its gain of toxicity, but on the other hand request very long NMR machine time and intense analysis, have not been defined in the previous report (9). Therefore, here we first targeted establishing a high-resolution picture of the conformations and dynamics of the co-existing two states of C71G-PFN1 by acquiring and analyzing a large set of different NMR experiments.

**Figure 2.**
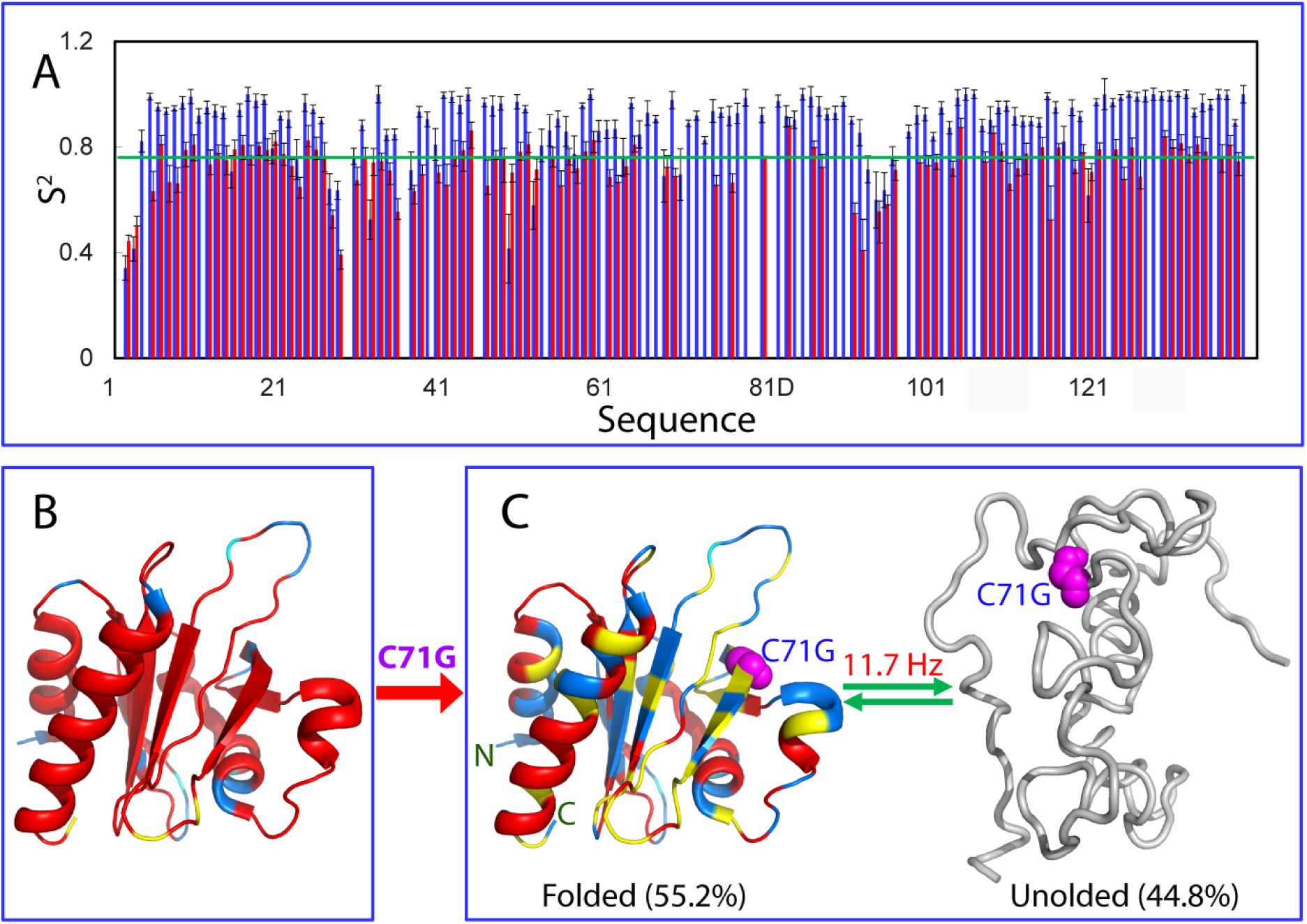
Conformations and dynamics of two coexisting states of C71G-PFN1. (A) Generalized squared order parameter (S^2^) of WT-PFN1 (blue) and the folded state of C71G-PFN1 (red). The green line has a S^2^ value of 0.76 (average – STD). (B) The structure of PFN1 (PDB ID of 2PAV) with S^2^ values of WT-PFN1 mapped on. (C). A diagram to show the co-existence of the folded and unfolded states of C71G-PFN1. S^2^ values are also mapped onto the folded state of C71G. Cyan is used for indicating Pro residues, and the yellow for residues with missing or overlapped HSQC peaks. Red is for residues with S^2^ values > 0.76 and blue for residues with S^2^ values < 0.76. The folded state with a population of 55.2% and unfolded state with a population of 44.8% have been experimentally mapped out to undergo a conformational exchange at 11.7 Hz.

We first collected triple resonance NMR spectra HN(CO)CACB and CBCA(CO)NH for achieving sequential assignments of both wild-type (WT) PFN1 and C71G-PFN1. As shown in Fig. S2, except for the C71G-PFN1 residues with missing or overlapped HSQC peaks, the (ΔCα-ΔCβ) values of all residues are highly similar for both WT-PFN1 and the folded state of C71G-PFN1 (Fig. S2A), indicating that the folded state of C71G-PFN1 has a conformation very similar to that of WT-PFN1. On the other hand, the absolute values of (ΔCα-ΔCβ) of the unfolded state are much smaller than those of the folded state of C71G-PFN1 (Fig. S2B), clearly suggesting that the unfolded state of C71G-PFN1 is highly disordered without any stable secondary and tertiary structures (32).

Subsequently, we acquired ^15^N backbone relaxation data T1, T2 and hNOE for both WT-PFN1 and C71G-PFN1 (Fig. S3), and performed the “model-free” analysis which generates squared generalized order parameters, S^2^, reflecting the conformational rigidity on ps-ns time scale. S^2^ values range from 0 for high internal motion to 1 for completely restricted motion in a molecular reference frame (31-35). As shown in Fig. 2A, the majority of the WT-PFN1 residues has S^2^ > 0.76 (average – STD), suggesting that WT-PFN1 has very high conformational rigidity over the whole structure (Fig. 1B) exactly as previously reported on WT-PFN1 (36). By contrast, many residues of the folded state of C71G-PFN1 have S^2^ < 0.76 (Fig. 2A), revealing that even the folded state of C71G-PFN1 becomes more dynamic than WT-PFN1 on ps-ns time scale (Fig. 2C). Interestingly, the overall rotational correlation time (τ_c_) of WT-PFN1 was determined to be 7.5 ns while that of C71G-PFN1 was 7.8 ns, implying that C71G-PFN1 becomes slightly less compact, completely consistent with the result that the S^2^ values of many residues of C71G-PFN1 becomes smaller than the corresponding ones of WT-PFN1.

Moreover, C71G-PFN1 has the co-existence of two sets of HSQC peaks (9): one from the folded and another from the unfolded state (Fig. S1A). Similar T2 (Fig. S3C) but very different T1 (Fig. S3B) values for most residues of two states imply that the folded and unfolded states of C71G-PFN1 undergo conformational exchanges mainly via longitudinal magnetization transfer (37). Therefore we collected HSQC-NOESY and indeed identified that the backbone amide protons of two states have cross-peaks as exemplified in Fig. S4. Subsequently, we selected a list of residues with well-separated NOE peaks and determined as previously described (31,37) the populations of the folded and unfolded states to be 55.2% and 44.8% respectively, while two states undergo conformational exchanges at 11.7 Hz (Table S1 and Fig. 2C).

We further assessed their hydrodynamic properties with pulsed field gradient NMR self-diffusion measurements on both WT-PFN1 and C71G-PFN1 as we previously performed on the TDP-43 N-terminal domain (31,38). The diffusion coefficient of WT-PFN1 was determined to be ~1.12 ± 0.03 × 10^−10^ m^2^/s, which is a typical value for a well-folded protein of 15 kDa (39). For C71G-PFN1, by fitting the very up-field NMR peaks only from the folded state, we first determined the diffusion coefficient of the folded state to be ~1.03 ± 0.02 × 10^−10^ m^2^/s, suggesting that C71G-PFN1 has a slight slower translational diffusion most likely due to a less compact packing than that of WT-PFN1 as reflected by S^2^ (Fig. 2A). Subsequently we also determined the diffusion coefficient of the unfolded state of C71G-PFN1 to be ~0.96 ± 0.02 × 10^−10^ m^2^/s, similar to that of chymotrypsinogen, a well-folded protein with a molecular weight of 25.6 kDa (39), indicating that the unfold state has much slower translational diffusion because of being disordered.

### NMR characterization of the interaction of C71G-PFN1 with FUS (1-267)

We titrated ^15^N-labeled C71G-PFN1 at 50 μM by step-wise addition of unlabeled FUS (1-267) as monitored by NMR 1D (Fig. 3A) and HSQC (Fig. 3B) spectra. As FUS (1-267) contains no large hydrophobic residues, it has no NMR 1D peaks < 1 ppm which are from the methyl groups of Val, Ile or Leu. On the other hand, the very up-field 1D peaks < 0.6 ppm are from only the folded state of C71G-PFN1.

**Figure 3.**
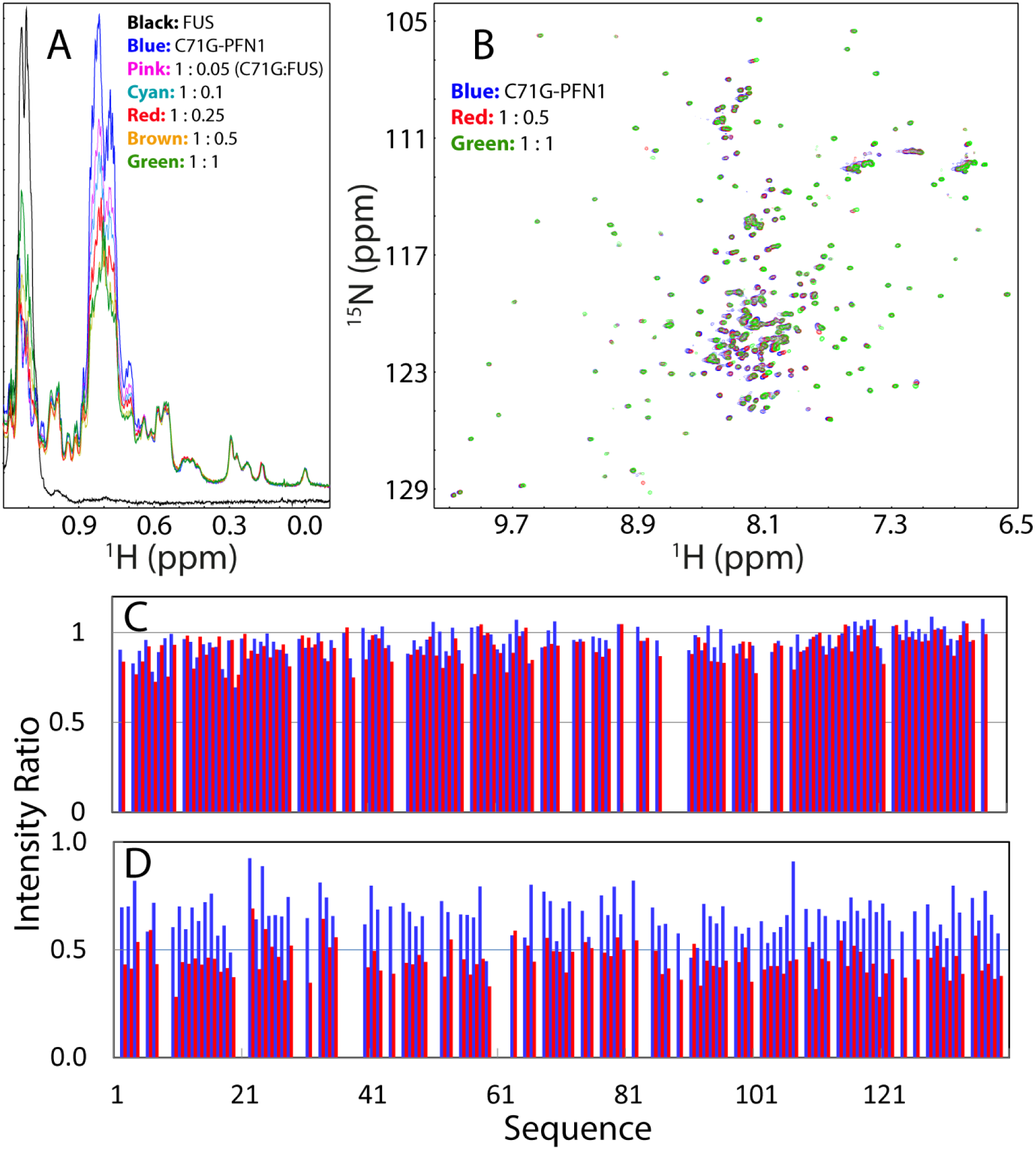
NMR characterization of ^15^N-labeled C71G-PFN1 interacting with unlabeled FUS (1-267). (A) Up-field 1D NMR spectra of C71G-PFN1 at 50 μM in the presence of FUS (1-267) at different molar ratios. As FUS (1-267) contains no large hydrophobic residues Leu, Ile and Val, it has no NMR peaks with chemical shifts < 0.9 ppm. As such, NMR peaks with chemical shifts < 0.5 ppm are from the folded state of C71G, while peaks with chemical shifts between 0.5 and 0.9 ppm are from both folded and unfolded states of C71G. (B) Superimposition of HSQC spectra of ^15^N-labeled C71G-PFN1 in the presence of unlabeled FUS (1-267) at different molar ratios. Residue-specific ratios of HSQC peak intensity of the folded (C) and unfolded (D) states of ^15^N-labeled C71G-PFN1 in the presence of unlabeled FUS (1-267) at molar ratio of 1:0.1 (blue) and 1:1 (red) (C71G:FUS).

Interestingly, upon gradual addition of FUS (1-267), both intensity and chemical shift of the very up-field peaks from the folded state of C71G-PFN1 remained almost unaffected, while the intensity of peaks over ~0.7-0.88 ppm from the methyl groups of Val, Ile or Leu of both folded and unfolded states reduced in a step-wise manner (Fig. 3A). Further analysis of HSQC spectra revealed that both chemical shift (Fig. 3B) and intensity (Fig. 3C) of HSQC peaks of the folded state of C71G-PFN1 were not significantly affected. By contrast, almost all HSQC peaks of the unfolded state were shifted (Fig. 3B), and intensity of most HSQC peaks reduced significantly even at a molar ratio of 1:0.1 (C71G:FUS); while at 1:1 the intensity of most HSQC peaks only account for ~40% (Fig. 3D). This strongly suggests that in the presence of FUS (1-267), the folded state was not significantly affected as evidenced by NMR results but the unfolded state appeared to acquire high dynamics on μs-ms time scale or/and become associated, and consequently its NMR peaks become significantly broadened. As FUS (1-267) at 50 μM could form liquid droplets in the presence of C71G-PFN1 at 1:1 (FUS:C71G) (Fig. 1D), which also gave high-quality NMR spectra, all further NMR studies of the complex between FUS (1-267) and C71G-PFN1 have been conducted under this condition.

We also conducted NMR self-diffusion measurements on FUS (1-267) alone at 50 μM and in the presence of C71G-PFN1 at a molar ratio of 1:1 (FUS:C71G). Due to being intrinsically disordered, FUS (1-267) alone has a diffusion coefficient of ~0.60 ± 0.01 × 10^−10^ m^2^/s, suggesting that it has much larger hydrodynamic radius and slower translational diffusion even than the unfolded state of C71G-PFN1. Interestingly, in the mixture, the diffusion coefficient of the folded state of C71G-PFN1 remained almost the same: ~1.04 ± 0.04 × 10^−10^ m^2^/s, while the diffusion coefficient of the unfolded state becomes ~0.81 ± 0.04 × 10^−10^ m^2^/s, similar to that of dimeric β-lactoglobulin with a molecular weight of 36.8 kDa (39). This result suggests that the translational diffusion of the unfold state of C71G-PFN1 becomes much slower due to the non-specific interaction with liquid droplets formed by FUS (1-267). Interestingly, the diffusion coefficient of FUS (1-267) in the mixture was determined to be 0.39 ± 0.03 × 10^−10^ m^2^/s, revealing that the translational diffusion of FUS was also significantly slowed down in the presence of C71G-PFN1.

### NMR dynamics of FUS (1-267) in the absence and in the presence of C71G-PFN1

Previously, FUS (1-165) has been extensively characterized by NMR spectroscopy and the study revealed that it remained to be highly disordered even in droplets by analyzing both NMR chemical shifts and relaxation data (26). Here we found that HSQC peaks of the isolated FUS (1-165) are almost completely superimposable to those of the corresponding residues in the context of FUS (1-267) (Fig. 5A). However, due to its extremely Gly-rich (54.9%) and highly degenerative sequence, the sequential assignment of the extra region over 166-267 of FUS (1-267) was not possible. So in our analysis we divided the unassigned residues over 166-267 into two categories: Gly residues with their HSQC peaks well-separated from the rest and non-Gly other residues. We titrated ^15^N-labeled FUS (1-267) at 50 μM with the unlabeled C71G-PFN1 and interestingly no detectable shift was observed for HSQC peaks of backbone amide protons up to the ratio of 1:1 (FUS:C71G) (Fig. 4B), implying that no specific interaction exists between FUS (1-267) and C71G-PFN1, completely consistent with the results obtained by monitoring HSQC peak intensity of the 15N-labeled C71G-PFN1 (Fig. 3D). Unexpectedly, however, detailed analysis showed a significant increase of the intensity of HSQC peaks for almost all FUS (1-267) residues at the ratio of 1:1 (FUS:C71G) (Fig. 4C).

**Figure 4.**
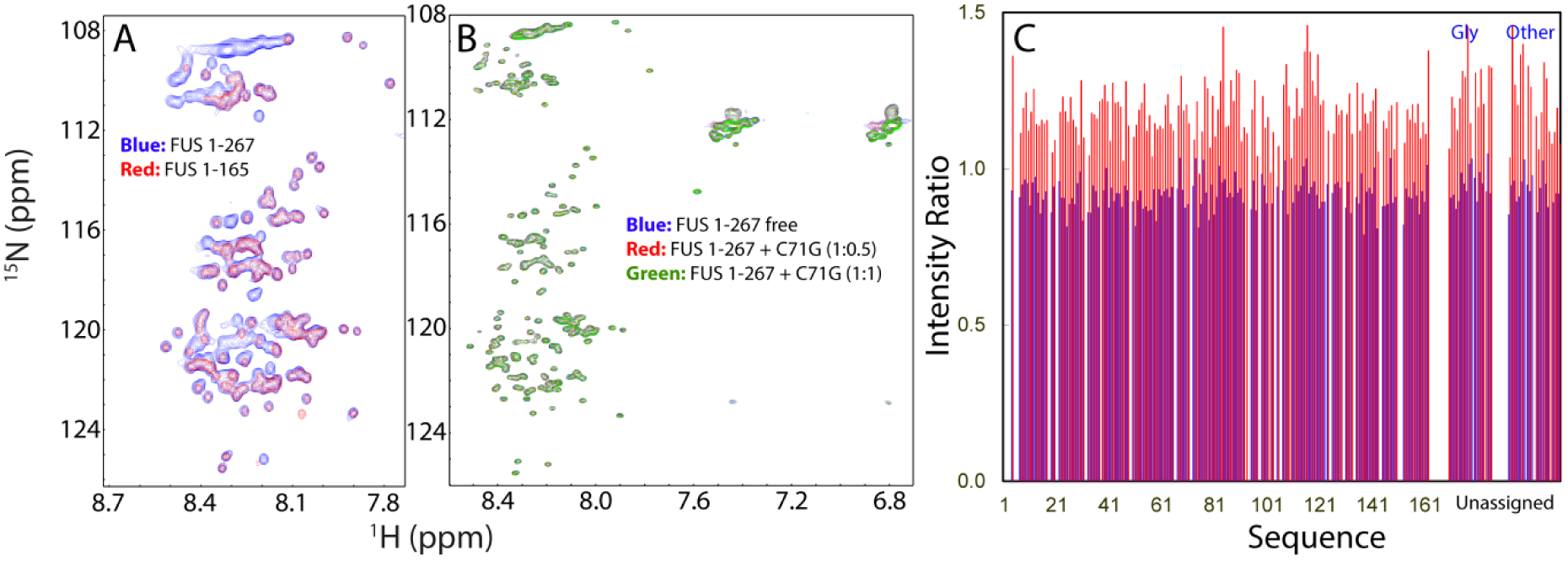
NMR characterization of ^15^N-labeled FUS (1-267) interacting with unlabeled C71G-PFN1. (A) Superimposition of HSQC spectra of ^15^N-labeled FUS (1-267) (blue) and FUS (1-165) (red). (B) Superimposition of HSQC spectra of ^15^N-labeled FUS (1-267) at 50 μM in the presence of unlabeled C71G-PFN1 at different molar ratios. (C) Residue-specific ratios of HSQC peak intensity of FUS (1-267) in the presence of unlabeled C71G-PFN1 at molar ratio of 1:0.1 (Blue) and 1:1 (red) (FUS:C71G).

**Figure 5.**
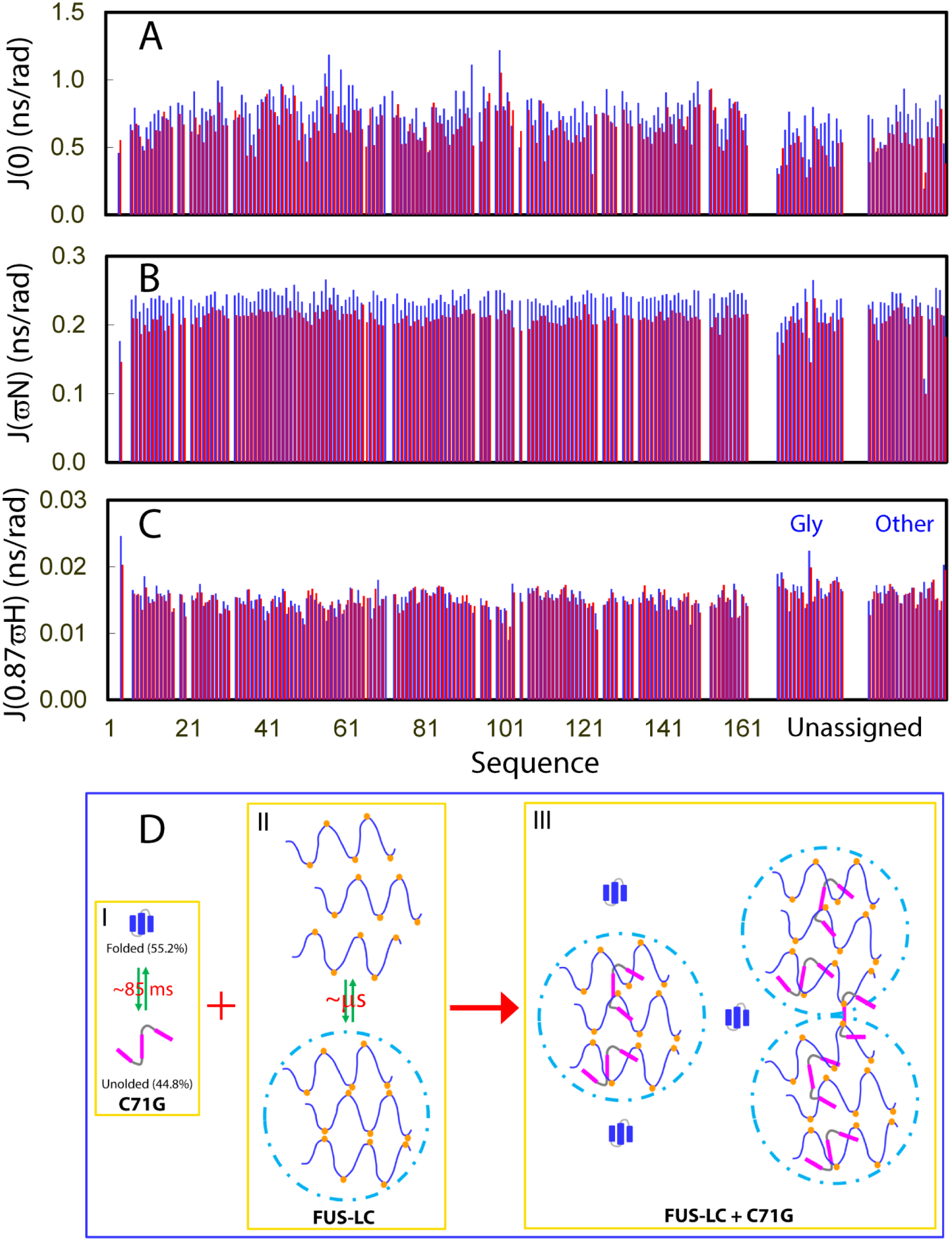
NMR dynamic views of FUS (1-267) interacting with C71G-PFN1. Spectral densities of ^15^N-labeled FUS (1-267) at 50 μM (blue); and in the presence of unlabeled C71G-PFN1 at a molar ratio of 1:1 (FUS:C71G), calculated from the ^15^N backbone relaxation data measured at 800 MHz. (A) J(0), (B) J(ωN), and (C) J(0.87ωH). (D) Proposed model by which the dynamic liquid droplets non-specifically and statistically trap/capture the unfolded state, but not the folded state of C71G-PFN1.

To understand this observation, we acquired the ^15^N backbone relaxation data T1, T2 and hNOE (Fig. S4), as well as Carr-Purcell-Meiboom-Gill (CPMG) relaxation dispersion (data not shown) data for FUS (1-267) in the absence and in the presence of C71G-PFN1 at 1:1 (FUS:C71G) under the solution condition exactly the same as that used for the DIC microscopy imaging (Fig. 1D). Previously the conformations of FUS (1-165) in solution and droplets have been characterized to be highly similar as well as to undergo exchanges faster than millisecond (25). Here, as indicated by the relaxation data (Fig. S5), the residues 1-165 are also highly disordered in the context of FUS (1-267), very similar to what was previously observed for the isolated FUS (1-165) (25). Moreover, despite being not assigned, the NMR relaxation data also provided critical insights into the dynamics of the extra Gly-rich region 166-271: both Gly and other non-Gly residues are also highly disordered. Furthermore, our CPMG data also showed that all resolved peaks have no significant dispersion (> 2 Hz) over the range from 40 Hz to 980 Hz, suggesting that as previously reported on FUS (1-165) (25), for FUS (1-267), the monomeric state and state in droplets are also undergoing conformational exchanges faster than millisecond. Very interestingly, in the presence of C71G-PFN1, most residues have increased T2 values (Fig. S5C), thus explaining the intensity increase of FUS (1-267) in the presence of C71G-PFN1 (Fig. 4C). This observation strongly implies that the presence of C71G-PFN1 might render FUS (1-267) to become more dynamic on ps-ns time scale, or/and reduces the conformational exchanges between the monomeric state and state in droplets. Furthermore, no CPMG dispersion > 2 Hz was observed also for FUS (1-267) in the presence of C71G-PFN1 at 1:1.

We then calculated reduced spectral densities at three frequencies, 0, ωN and 0.87ωH (Fig. 5) from the ^15^N backbone relaxation data, which reflect relaxation contributions from motions/exchanges on different timescales (32, 40-41). Generally, fast internal motions on ps-ns timescale are expected to reduce the value of J(0), while slow motions/exchanges on μm-ms time scale lead to large values of J(0). On the other hand, the high-frequency spectral density J(0.87ωH) is only sensitive to fast internal motions, which result in relatively large values of J(0.87ωH). Interestingly, in the presence of C71G-PFN1, most residues of FUS (1-267) have reduced J(0) values (Fig. 5A), indicating the increase of internal motions on ps-ns timescale, or/and the decrease of slow motions/exchanges on μm-ms time scale. However, as no significant change of J(0.87ωH) values was observed for FUS (1-267) in the free state and in the presence of C71G-PFN1 (Fig. 5C), this result indicates no significant increase of internal motions of FUS (1-267) on ps-ns timescale in the presence of C71G-PFN1. Taken together, the results imply that the accumulation of C71G-PFN1 within the droplets might affect the exchange processes between the monomeric state and state in droplets. In the future, it is of fundamental interest to better understand this observation despite the extreme challenge.

## Discussion

Although whether common mechanisms exist for their gain of toxicity remain elusive, emerging evidence implies that misfolded proteins are toxic in general and their chaperone-assisted refolding or degradation not only represents a central task for cells to maintain proteostasis, but also a key step to rejuvenate. Indeed, the naked mole rat apparently without aging (42) can maintain its proteostasis exceptionally good by minimizing misfolded proteins with increased translational fidelity (43), high resistance of proteins to oxidative damages (44); and enhanced activity of the degradation machinery (45). For a PQC machinery to precisely remove misfolded proteins, a pivotal component required is an efficient mechanism to recognize and recruit misfolded proteins. Recent studies indicate that SGs act as a surveillance system, which can efficiently recruit misfolded proteins accumulated during cellular stress by a non-classic mechanism with dynamic liquid droplets formed by LC domains of SG components such as FUS and TDP-43 (20-22). On the other hand, however, the aberrant dynamics of liquid droplets resulting from accumulation of misfolded proteins may in fact represent a mechanism of gain of toxicity for misfolded proteins, as also implicated by the increasing observations that misfolded proteins generally disrupted the dynamics of SGs (1,10-15,20-29). Therefore, to delineate this mechanism is not only of central importance to understanding fundamental cellular physiology, but also offers a key to developing therapeutic strategies to treat protein-aggregation diseases or even aging.

So far, the molecular mechanism for liquid droplets to recruit misfolded proteins still remains undefined, and in this regard biophysical characterizations particularly by NMR spectroscopy can provide high-resolution mechanistic insights. In the present study, we first assessed the effects of three ALS-causing PFN1 mutants on dynamics of liquid droplets of FUS (1-267). The results indicate for the first time that without needing any other factors, ALS-causing PFN1 mutants are sufficient to disrupt dynamics of liquid droplets of FUS (1-267) with an order of the disrupting capacity as: C71G > G118V > E117G, which is amazingly correlated to the ALS-causing toxicity of three mutants (3,6,8,46). Noticeably, their disrupting capacity also appears to be overall correlated with their populations of the unfolded states (Fig. S1), as well as their tendency in forming aggregation/amyloid fibrils in cells and *in vitro* (3,6,46). This observation thus implies that likely the unfolded state of PFN1 accounts for the disrupting capacity for liquid droplets. Indeed, TDP-43 (10-102) with only the unfolded state seemingly has the disrupting capacity higher than PFN1 mutants.

Subsequently, by acquiring and analyzing a variety of multi-dimensional NMR experiments we characterized conformations, dynamics and hydrodynamic properties of C71G-PFN1, FUS (1-267) and their interaction. The results reveal that while the solution conformation of the folded state of C71G-PFN1 is very similar to that of WT-PFN1, the unfolded state is predominantly disordered without any stable secondary and tertiary structures. On the other hand, however, even the folded state of C71G-PFN1 becomes more dynamic than WT-PFN1 on ps-ns time scale. Furthermore, by quantitatively analyzing NMR spectra reflecting chemical exchange process via longitudinal magnetization transfer, the populations of the folded and unfolded state were determined to be ~55.2% and ~44.8% respectively, which undergo conformational exchange at 11.7 Hz. As such, C71G-PFN1 represents an excellent model for misfolded proteins, which offers unique advantage for characterizing the interactions of liquid droplets with both folded and misfolded states under the exact same conditions by the same sets of NMR experiments. As out of three ALS-causing PFN1 mutants studied here, the most toxic C71G-PFN1 has the highest capacity in disrupting dynamics of liquid droplets formed by FUS (1-267), we further performed extensive NMR characterizations of the interaction between C71G-PFN1 and liquid droplets formed by FUS (1-267). The results reveal that while the folded state of C71G-PFN1 is not significantly affected by liquid droplets, the unfolded state strongly interacts with liquid droplets in a non-specific manner.

Our results all together thus allow the proposal of a model for the liquid droplets formed by the LC domains to recruit misfolded proteins (Fig. 5D). As the conformational exchange between the folded and unfolded states of C71G-PFN1 occurs at ~85 ms (I of Fig. 5D), much slower than the μs dynamic equilibrium between the monomeric state and state in droplets of FUS (1-267) (II of Fig. 5D), the liquid droplets are thus able to differentiate the folded and unfolded states of C71G-PFN1. For the folded state of C71G-PFN1 with a tight packing and globular shape, if its size is larger than the interior space of liquid droplets it will be excluded from entering the droplets. If its size is smaller than the interior space of liquid droplets it may enter the droplets but statistically behaves as being able to freely pass through the droplet. Overall, the folded state appears not to be significantly affected by the droplets. By contrast, as the unfolded state of C71G-PFN1 is an ensemble of extended conformations exchanging on ps-ns time scale without any stable secondary and tertiary structures, even without needing specific interactions with the droplets, it can be statistically trapped/captured within the liquid droplets (III of Fig. 5D). Moreover, some unfolded C71G-PFN1 molecules might be trapped within two droplets (III of Fig. 5D), thus leading to the observation that C71G-PFN1 is able to trigger clustering/coalescing of the different droplets (Fig. 2C and 2D; Video 2C and 2D). This also explains the amazing observation that TDP-43 (10-102) with only the unfolded conformation can even connect many droplets in the “beads-in-a-string” style (Fig. 1I-1J and Video 1I-1J).

Nevertheless, this mechanism of recruiting misfolded proteins appears to function at great cost: the dynamics of the droplets will be reduced even simply due to accumulation of unfolded molecules within the droplets. Moreover, under certain stress conditions, misfolded proteins are largely generated, and thus not all of them can be efficiently refolded by chaperones. In particular, for patients carrying disease-causing genetic variations, the unfolded states of these mutants are no longer refoldable due to the loss of their intrinsic capacity to completely fold as exemplified by L126Z-SOD1 and C71G-PFN1 (47,48). As a result, these misfolded/unfolded proteins will become significantly accumulated within SGs. If they also cannot be rapidly degraded, they might become irreversibly co-precipitated with other SG components driven by non-specific hydrophobic interactions between the recruited misfolded proteins and the exposed hydrophobic regions of SG proteins as previously reported on TDP-43 (15,16); or/and self-associated and aggregated which can be significantly accelerated by the high local concentrations of misfolded proteins within the liquid droplets because all misfolded/unfolded proteins are aggregation-prone (4,8,46-48). As a consequence, the structures, dynamics and functions of SGs might be disrupted or even destroyed by misfolded proteins, and this will manifest as gain of toxicity for misfolded proteins.

In summary, our present study provides mechanistic insights into the mechanism by which dynamic liquid droplets formed by the LC domains to recruit misfolded proteins. In light of this mechanism, ALS-causing PFN1 mutants might gain toxicity to trigger ALS by disrupting the dynamics of SGs organized by LLPS, thus strongly supporting the emerging observation that disruption of the dynamics of SGs by disease-associated proteins/mutants represents a key mechanism underlying various neurodegenerative diseases including ALS (1,10-15,20-29). In particular, ALS-causing PFN1 mutants C71G-, G118V and E117G-PFN1 are fundamentally indistinguishable from other misfolded/aggregation-prone proteins as evidenced by the result that indeed unfolded TDP-43 (10-102) is also able to disrupt LLPS. On ther other hand, LLPS representing a common principle to organize cellular membrane-less organelles lies at the heart of many essential cellular functions which even include the formation of heterochromatin domains and cell cycle regulation (27, 49-52). Therefore, misfolded proteins might share a common capacity in disrupting LLPS for gain of toxicity.

## Methods

### Preparation of protein samples

The expression and purification of WT-PFN1, C71G-, G118V- and E117G-PFN1, as well as removal of His-tag followed the protocol we previously described (9). For FUS (1-267), here to avoid the perturbation of His-tag in liquid-liquid phase separation (LLPS), a stop codon was added immediately after the DNA sequence encoding FUS (1-267) in a modified vector pET28a with a C-terminal His-tag, we previously used for expressing His-tagged FUS (1-267) (28). Consequently, the present FUS (1-267) recombinant protein was purified exactly as previously described (28) except for that the Ni^2+^-affinity purification is no longer needed. In the present study, DTT was added into all samples containing WT-PFN1 or PFN1 mutants, or TDP-43 (10-102) to a final concentration of 1 mM to prevent the non-native cross-linkage of free Cys residues.

The generation of the isotope-labeled proteins for NMR studies followed a similar procedure except that the bacteria were grown in M9 medium with the addition of (^15^NH_4_)_2_SO_4_ for ^15^N labeling and (^15^NH_4_)_2_SO_4_/[^13^C]-glucose for double labelling (9,28). The purity of the recombinant proteins was checked by SDS–PAGE gels and their molecular weights were verified by a Voyager STR matrix-assisted laser desorption ionization time-of-flight-mass spectrometer (Applied Biosystems). The concentration of protein samples was determined by the UV spectroscopic method in the presence of 8 M urea (53).

### Differential interference contrast (DIC) microscopy

Characterization of liquid droplets was conducted for different samples at a protein concentration of 50 μM in 1 mM sodium phosphate buffer (pH 6.0) at 25 °C, which was imaged as previously described (30) by differential interference contrast (DIC) microscopy (OLYMPUS IX73 Inverted Microscope System with OLYMPUS DP74 Color Camera).

### NMR experiments

All NMR experiments were acquired at 25 °C on an 800 MHz Bruker Avance spectrometer equipped with pulse field gradient units as described previously (9,28). NMR data were processed with NMRPipe (54) and analysed with NMRView (55). For achieving sequential assignments of WT-PFN1 and C71G-PFN1, NMR experiments including HNCACB and CBCA(CO)NH were acquired on ^15^N-/^13^C-double labelled samples in 1 mM sodium phosphate buffer (pH 6.0). For other NMR studies on interactions between FUS (1-267) and C71G-PFN1, the NMR samples were prepared at protein concentration of 50 μM in 1 mM sodium phosphate buffer (pH 6.0).

### PGF Diffusion Measurement

PGF-NMR experiments were run on an 800 MHz Bruker Avance spectrometer at 25°C. The different protein samples were prepared in D_2_O at protein concentration of 50 μM in 1 mM sodium phosphate buffer (pD 6.0). The experiments were performed using the Bruker pulse sequence and the Bruker macro diffusion ordered spectroscopy (DOSY) (31,38). Typically 16 values of gradient strength were used in the range 0 to 32 G/cm, with PFG duration of 2 ms, and diffusion time of 150 ms. The self-diffusion coefficients (*D_s_*) were calculated using the Bruker DOSY analysis program. Each sample was run in triplicate and *Ds* values were averaged over the three experiments. The resulting decay curves were fitted and *Ds* values were calculated with the equation below:

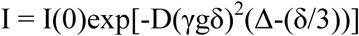

Where I(0) is 1.002, γ is 4.258×10^3^ Hz/G, δ is 4.000 ms, and Δ is 150 ms.

### NMR ^15^N backbone dynamics on ps-ns time scale

^15^N backbone T1 and T1ρ relaxation times and {^1^H}-^15^N steady state NOE intensities were collected on the ^15^N-labeled C71G-PFN1 as well as ^15^N-labeled FUS (1-267) in the presence and in the presence of unlabelled C71G-PFN1 at a molar ratio of 1:1 (FUS:C71G) at 25 °C at protein concentration of 50 *μ*M in 1 mM phosphate buffer (pH 6.0) on an Avance 800 MHz Bruker spectrometer with both an actively shielded cryoprobe and pulse field gradient units (28,34,35). Relaxation time T1 was determined by collecting 7 points with delays of 10, 160, 400, 500, 640, 800 and 1000 ms using a recycle delay of 1 s, with a repeat at 400 ms. Relaxation time T1*ρ* was measured by collecting 8 points with delays of 1, 40, 80, 120, 160, 200, 240 and 280 ms, with a repeat at 120 ms. {^1^H}-^15^N steady-state NOEs were obtained by recording spectra with and without ^1^H presaturation, a duration of 3 s and a relaxation delay of 6 s at 800 MHz.

### Model-free analysis

NMR relaxation data were analyzed by “Model-Free” formalism with protein dynamics software DYNAMICS (28, 32-35). Briefly, relaxation of protonated heteronuclei is dominated by the dipolar interaction with the directly attached ^1^H spin and by the chemical shift anisotropy mechanism. Relaxation parameters are given by:

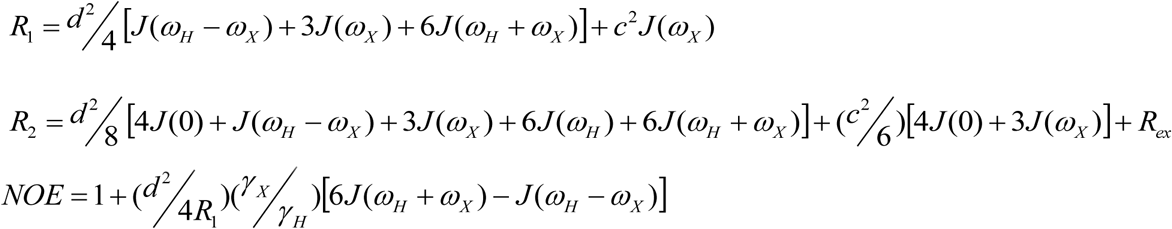

In which,

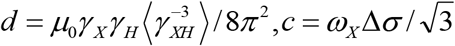 *μ*_0_ is the permeability of free space; *h* is Planck’s constant; *γ_X_*, *γ_H_* are the gyromagnetic ratios of ^1^H and the X spin (X=^13^C or ^15^N) respectively; *γ _XH_* is the X-H bond length; *ω_H_* and *ω_X_* are the Larmor frequencies of ^1^H and X spins, respectively; and Δ*σ* is the chemical shift anisotropy of the X spin.

The Model-Free formalism determines the amplitudes and time scales of the intramolecular motions by modeling the spectral density function, *J*(*ω*), as

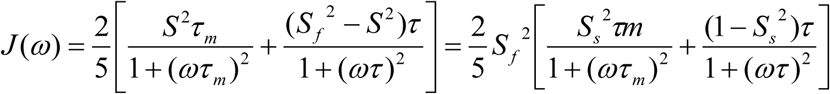

In which, 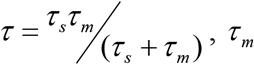 is the isotropic rotational correlation time of the molecule, *τ_s_* is the effective correlation time for internal motions, 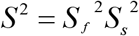 is the square of the generalized order parameter characterizing the amplitude of the internal motions, and 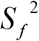 and 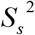 are the squares of the order parameters for the internal motions on the fast and slow time scales, respectively.

In order to allow for diverse protein dynamics, several forms of the spectral density function, based on various models of the local motion, were utilized, which include the original Lipari-Szabo approach, assuming fast local motion characterized by the parameters *S*^2^ and *τ_loc_*; extended model-free treatment, including both fast 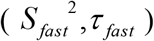 and slow 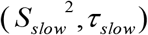 reorientations for the NH bond (*τ_fast_* << *τ_slow_* < *τ_c_*); and could also allow for slow, milli-to microsecond dynamics resulting in a conformational exchange contribution, *R_ex_*, to the linewidth. In DYNAMICS, there are eight models for local motions and each residue is fitted with different models. Subsequently goodness of fit will be checked and the best-fitted model will be selected.

The relaxation data of WT-PFN1 and the folded state of C71G-PFN1 were analyzed with the previously published X-ray structure (pdb ID of 2PAV) (56) by isotropic, axially-symmetric and fully anisotropic models for the overall motion and the results were tested and then compared. According to the illustration of ROTDIF, isotropic model was finally selected for both WT-PFN1 and the folded state of C71G-PFN1 because of smallest Ch^2^/df value. For WT-PFN1, τc = 7.5 ns and Dx = Dy = Dz = 1.852 E ± 07 s^-1^; for the folded state of C71G-PFN1, τc = 7.8 ns and Dx = Dy = Dz = 2.094 E ± 07 s^-1^.

### Reduced spectral density analysis

The ^15^N backbone relaxation data of FUS (1-267) in the absence and in the presence of unlabeled C71G-PFN1 were analyzed by direct mapping of the reduced spectral density with simplified approximations (40,41). Briefly, J(0), J(ωN), and J(0.87ωH), the spectral densities at the frequencies 0, ωN, and 0.87ωH respectively, were calculated based on Eqs.:

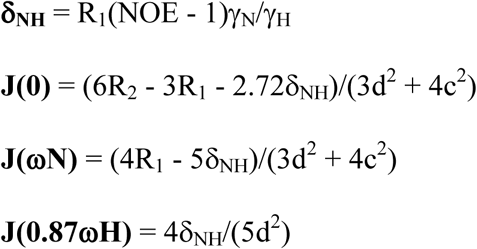

where d = (μ0hγ_N_ γ_H_/8π^2^) (r^-3^) and c = ωNΔδ/3^1/2^.

### NMR ^15^N backbone dynamics on μs-ms time scale

^15^N transverse relaxation dispersion experiments were acquired at 25 °C on the ^15^N-labeled FUS (1-267) in the absence and in the presence of unlabelled C71G-PFN1 at protein concentration of 50 μM in 1 mM phosphate buffer (pH 6.0) on a Bruker Avance 800 spectrometer. A constant time delay (*T*_CP_ = 50 ms) was used with a series of CPMG frequencies, ranging from 40 Hz, 80 Hz, 120 Hz (x2), 160 Hz, 200 Hz, 240 Hz, 320 Hz, 400 Hz, 480 Hz, 560 Hz, 640 Hz, 720 Hz, 800 Hz, and 960 Hz (x2 indicates repetition). A reference spectrum without the CPMG block was acquired to calculate the effective transverse relaxation rate by the following equation:

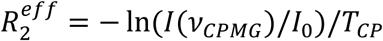

Where I(ν_CPMG_) is the peak intensity on the difference CPMG frequency, I_0_ is the peak intensity in the reference spectra (28,35).

### Quantification of Exchange Kinetics

Longitudinal magnetization transfer due to chemical exchange is the basis for the appearance of exchange cross peaks in nuclear overhauser effect spectroscopy (NOESY) spectra. The evolution of longitudinal magnetization is described by (37):

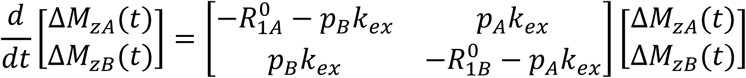

In which 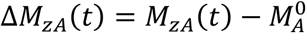 and 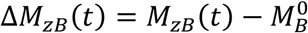. The solution of this equation is:

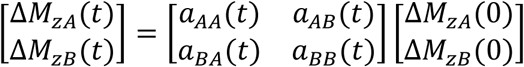

In which

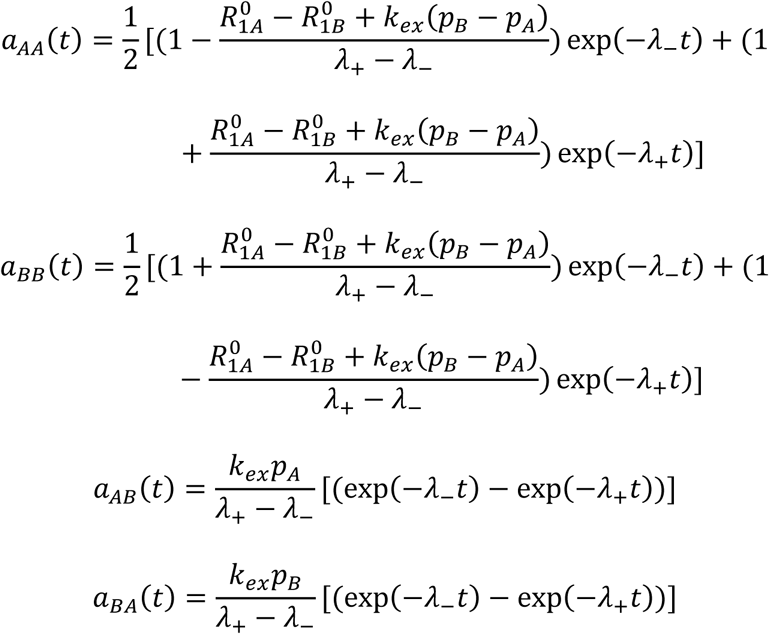

and

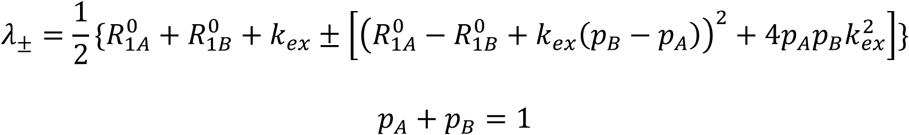

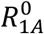 and 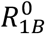 were measured by separate experiments. As a consequence, here *k_ex_*, *p_A_*, and *p_B_* could be derived as previously described (31,37) and presented in Table S1.

## Acknowledgments

This study is supported by Ministry of Education of Singapore Tier 2 Grant MOE2015-T2-1-111 (to J.S.).

## Author contributions

J.S. conceived and designed the research; J.K., L.-Z.L., and J.S. performed research; L.Z.L., J.K., and J.S. analyzed data; and J.S. wrote the paper.

